# Dual-Field Microvascular Segmentation: Hemodynamically-Consistent Attention Learning for Retinal Vasculature Mapping

**DOI:** 10.1101/2024.11.27.625635

**Authors:** Jingling Zhang, Xiangfei Liu, Shuting Zheng, Wen Zhang, Jia Gu

## Abstract

Accurate retinal Microvascular segmentation demands a balanced combination of anatomical fidelity and hemodynamic relevance. However, existing methods fall short in preserving critical structures such as capillary junctions and bifurcations, thus limiting clinical applications and causing fragmentation. To address these limitations, we propose DFMS-Net, a novel dual-field segmentation framework that synergistically integrates geometric-field modeling—achieved through the Spatial Pathway Extractor (SPE) and Transformer-based Topology Interaction (TTI) for preserving structural continuity—and functional-field optimization—realized by the Semantic Attention Amplification (SAA) module for enhancing semantic visibility—via a unified Dual-Field Hemodynamic Attention (DFHA) mechanism. This core module enables joint enhancement of vessel continuity, accurate resolution of complex branching patterns, and recovery of low-contrast capillaries, all under physiological guidance. By co-optimizing geometric and functional cues within a unified attention learning paradigm, DFMS-Net produces segmentations that are both morphologically accurate and hemodynamically plausible. Furthermore, we propose two specialized variants using a streamlined Double SPE Attention for vessel continuity refinement. To address the directionality-dependent nature of structural damage in retinal ischemia and glaucoma, we propose Variant-1—a streamlined architecture that emphasizes dual-stage directional refinement to enhance trajectory coherence and improve the detection of topological disruptions, while Variant-2 supports high-resolution analysis of capillary dropout in early diabetic retinopathy through detailed microvascular recovery. Extensive experiments on retinal (DRIVE, STARE) and coronary angiography (DCA1, CHUAC) datasets demonstrate that DFMS-Net achieves state-of-theart performance. Meanwhile, its strong generalization capability offers a promising foundation for diagnosing both retinal and cardiovascular diseases. The code will be available at https://github.com/699zjl/DFMS-Net-new.

## I. INTRODUCTION

PRECISE mapping of retinal vasculature networks, integrating both structural-spatial integrity and hemodynamically relevant mapping, forms a critical diagnostic cornerstone in the evaluation of multi-branched vessel disorders [1], [2]. The morphological characteristics of retinal vessels, such as vascular width, tortuosity, and branching patterns, which act as sensitive biomarkers for ocular pathologies, play a critical role in diagnosing and predicting the prognosis of common eye diseases like diabetic retinopathy, glaucoma, and age-related macular degeneration [1], [3]. Beyond ophthalmology, emerging evidences suggest that the retinal microvasculature also has far-reaching effects on the occurrence of circulatory system diseases. Retinal microvascular abnormalities can increase the risk of stroke by 2-to 3-fold and are closely connected to the generation of subclinical atherosclerosis [4]. Regarding hemodynamics, analyzing retinal vascular hemodynamics facilitates the early discovery of abnormalities in circulatory system-related diseases, including stroke, ischemic heart disease, heart failure, and renal dysfunction. Thus, clinicians can implement preventive strategies prior to the manifestation of detectable morphological alterations [2]. Prior studies demonstrate that medical image segmentation is the fundamental task of hemodynamic simulations, and the quality of segmentation results (e.g., the geometry of the vessel) directly affects hemodynamic simulation outcomes [5], [6], [7]. Considering the clinical significance of the structural spatial features and hemodynamically-informed analysis for fundus and blood-related diseases, developing automated retinal segmentation methods capable of tackling both geometric topology and blood flow consistency has become an urgent task.

However, automated retinal vessel segmentation remains a challenging task due to the inherent complexity (e.g., multi-scale vessels, overlapping structures), imaging limitations (low contrast, noise, motion artifacts), and the tree-like architecture with sub-resolution capillaries [3], [8]. Moreover, the limited scale of medical image datasets presents major challenges for effective model training [9], [10], [11]. While deep learning-based methods have significantly advanced retinal vessel segmentation, they are inherently short of the ability to maintain both structural continuity and hemodynamic coherence. Attention-based models (e.g., ECANet [12], Attention U-Net [13] and CBAM [14] lack directional priors for vascular trajectory modeling, resulting in fragmented predictions. In contrast, topology-aware methods,including Discrete Morse Theory [15], clDice [16], and centerline boundary-aware losses [17], aim to preserve vascular connectivity, while TopoMortar [18] provides a benchmark for topological evaluation. Yet all of them rely either on handcrafted rules or loss-level constraints, failing to embed hemodynamic consistency into feature learning processes.

To address this limitation, we propose DFMS-Net, a dual-field microvascular segmentation framework in which geometric and functional modeling are synergistically integrated through a Dual-Field Hemodynamic Attention (DFHA). Our key contributions are as follows:

- We propose DFMS-Net, a novel dual-field microvascular segmentation framework that integrates geometric-field modeling and functional-field optimization through a Dual-Field Hemodynamic Attention (DFHA) mechanism. The geometric field is jointly modeled by the Spatial Pathway Extractor (SPE) and Transformer-based Topology Interaction (TTI) to preserve vessel continuity and resolve complex branching patterns, while the functional field is enhanced via the Semantic Attention Amplification (SAA) module to recover faint capillaries under physiological guidance.
- We design the The Spatial Pathway Extractor (SPE) that employs axial-directional attention to maintain vessel trajectory integrity, reducing fragmentation in fine-scale microvasculature.
- We introduce the Transformer-based Topology Interaction (TTI) module,which captures long-range structural dependencies by modeling interactions between bifurcation points via learnable, physiology-informed adjacency matrices, resolving intricate branching patterns in anatomically plausible configurations.
- We develop the Semantic Attention Amplification (SAA) module to enhance local intensity changes of low-contrast capillaries and suppress background noise of medical images.

## II. RELATED WORK

### A. Deep Learning for Medical Segmentation

Deep Learning based architectures have established foundational standards in medical image analysis, with U-Net [19] remaining the cornerstone. Its encoder-decoder design effectively balances the preservation of local details and the integration of global context. Numerous variants have been proposed, including FR-UNet [20] for restoring the lost spatial information, I^2^U-Net [21] for bridging the encoder and decoder semantic gap, and MSVM-UNet [22] for comprehensive contextual integration across vascular hierarchies. Attention mechanisms have further enhanced feature discriminability: channel attention [12], [23] recalibrates feature importance, while spatial attention [24] improves localization under anatomical variability. More recently, transformer-based models (e.g., TransUNet [25], MISSFormer [26]), and generative models (e.g., MedSegDiff [27]) for noise-robust segmentation, and graph-based approaches (e.g., DPL-GFT-EFA [28]) for topological reasoning. Despite these advances, most methods treat segmentation as a pixel-wise task, often failing to preserve vascular continuity across bifurcations.

### B. Large Vision Models for Medical Segmentation

While prompts enable flexible interaction in vision models, their dependence on manual annotation limits clinical applicability in medical imaging. To alleviate this burden, recent works progress toward automation: a) *prompt-dependent* methods, starting from the original SAM [29], require explicit user input; Extensions like MedSAM [30], MedLSAM [31], and Samus [32] adapt SAM to medical data but still rely on point or box prompts; b) *semi-automatic* approaches generate prompts via learned priors—e.g., MoSE [33] uses a shape dictionary and PGP-SAM [34] leverages prototypes; c) *prompt-free* inference is achieved by MedSAMix [35] via model merging, and MedicOSAM [36] aims for a unified foundation model. Despite this evolution, these methods treat segmentation as a geometric recovery task, ignoring hemodynamic continuity and vascular topology,which are essential for accurate microvascular analysis.

### C. Topology-Aware Medical Segmentation

Accurate microvascular analysis relies heavily on maintaining vessel connectivity. To achieve this, Skeleton-based losses—including clDice [16], centerline-Dice [17], and skeleton recall loss [37]—are used to improve topological integrity through centerline alignment. Region-wise modeling [38] and skeleton-driven boundary refinement [39] further improve reconstruction of local structures. Some approaches utilize Discrete Morse Theory to achieve theoretically grounded topological control [15], whereas TopoMortar [18] offers a specialized benchmark for topology evaluation. Despite these improvements, recent methods tend to emphasize geometric consistency while neglecting the semantic context that is essential for robust connectivity prediction in morphologically complex or low-signal areas.

## III. METHODOLOGY

### A. Network Overview

Fig. 1 presents the dual-field microvascular segmentation Network (DFMS-Net), a hemodynamically-aware inspired architecture that revolutionizes retinal vasculature mapping through synergistic geometric-functional learning. Building upon baseline FR-UNet [20], our framework consists of three main components: the encoder module, the hemodynamic attention mechanism and the decoder module.

**Fig. 1.**
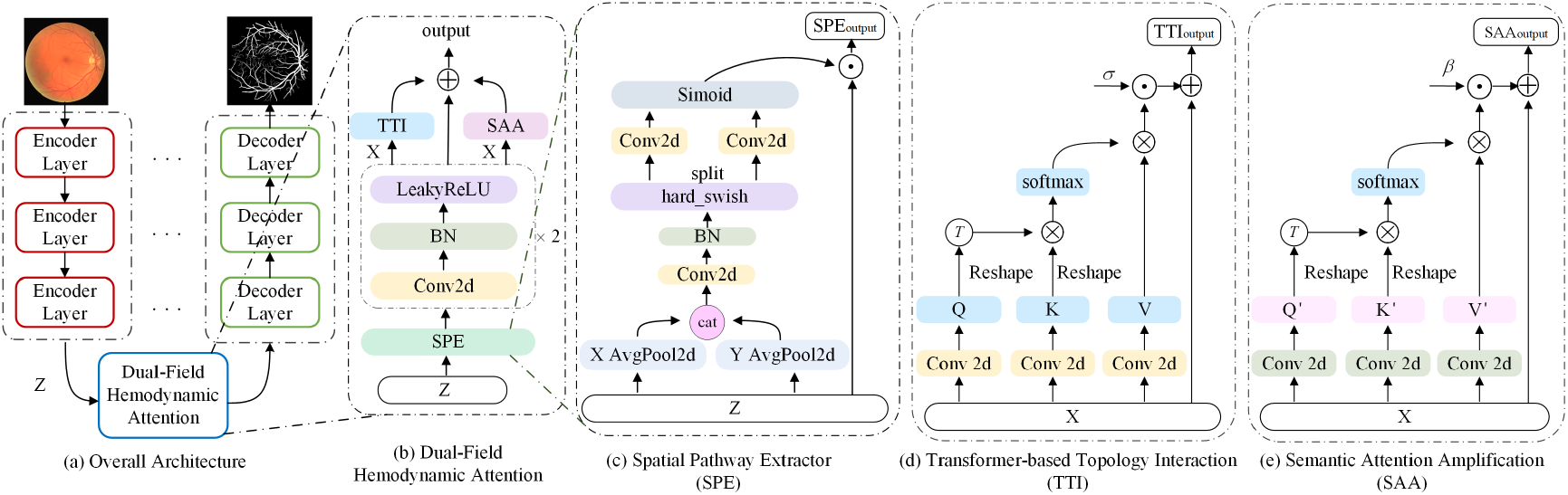
Architecture overview of the proposed DFMS-Net. The framework consists of three main components: the encoder module, the Dual-Field Hemodynamic Attention (DFHA) module, and the decoder module. The DFHA integrates three complementary components within the bottleneck layer: SPE and TTI for geometric-field modeling, and the SAA for functional-field optimization.

The encoder and decoder modules contain three stages of down-sampling and up-sampling layers, implemented by the standard 2 × 2 convolution and the 2×2 fractionally-strided convolutions, respectively. Each stage of the encoder and decoder modules comprises multiple convolution blocks. Unlike the FR-Unet, we substitute the modified residual block with a newly-designed standard convolutional block, which consists of two 3×3 convolution (Conv) layers, two batch normalization (BN) layers and two LeakyReLU (LR) activation layers. To generate dense predictions, we retain the skip-connection structure, which fuses the feature maps from the previous block with those obtained from the down-sampling and upsampling layers of adjacent stages.

Our core innovation lies in the Dual-Field Hemodynamic Attention (DFHA), which integrate three complementary components: Spatial Pathway Extractor (SPE), Transformer-based Topology Interaction (TTI), and Semantic Attention Amplification (SAA). These components work synergistically to address dual-field requirements—geometric-field structural modeling and functional-field semantic optimization—under physiological guidance. We now elaborate on each component in detail below.

### B. Dual-Field Hemodynamic Attention

This section details the design and operation of the three key components within the Dual-Field Hemodynamic Attention, guided by principles of blood flow dynamics and vascular morphology.

#### 1) Spatial Pathway Extractor (SPE)

Retinal microvasculature exhibits delicate capillary networks with highly variable tree-like topologies, where preserving vascular trajectory continuity is essential for accurate segmentation. However, conventional methods often fail to maintain long-range spatial coherence due to noise, low contrast, or abrupt intensity changes, leading to fragmented vessel paths and erroneous connections.

To address this challenge, we introduce SPE, which leverages hemodynamic principles to model directional flow patterns along potential vessel pathways. Specifically, SPE employs an axial-awareness attention mechanism that simulates blood flow directionality by aggregating features along dominant vessel orientations—horizontal and vertical directions—thereby enhancing spatial coherence and suppressing false connections.

Given a feature map *Z* ∈ ℝ^*C×H×W*^, which is directly generated from the third layer of the encoder module. Here *H, W* and *C* denotes the height, width and channel number of the feature map, respectively. The directional feature maps are then embeded by two orthogonal average poling with kernel sizes of (*H*,1) and (1,*W*), dependently [40]. These pooling operations simulate the clinical practice of tracing vessel continuity along their dominant orientations. Horizontal pooling (kernel (*H*,1)) aggregates intensity profiles across the vessel’s width, suppressing noise while preserving its longitudinal continuity:

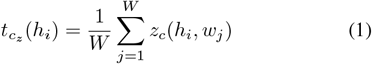

Conversely, vertical pooling (kernel (1,*W*)) captures cross-sectional variations critical for detecting bifurcation points:

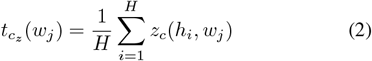

Here, *h*_*i*_ (*i* ∈ [1, *H*]) denotes the height index, *w*_*j*_ (*j* ∈ [1, *W*]) denotes the width index, and *c*_*z*_ (*z* ∈ [1, *C*]) denotes the channel index. The transformed features of the *c*_*z*_ channel at each spatial location index (*h*_*i*_, *w*_*j*_) fuse the long-rang direction-sensitive relationships and spatial location information, which are critical to model complex strcucture details. The pooled 1D features encode global vascular geometry, but may overlook local texture details (e.g., capillary wall irregularities). To reconcile global continuity with local fidelity, those two orthogonal embeddings are concatenated and then projected into a composite signal *p*_*c*_ ∈ ℝ^2*H×*1^ through a 1 × 1 convolutional filter *f* ^1*×*1^, which acts as a differentiable vessel edge detector:

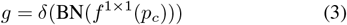

Here, the hard-swish activation function (*δ*) is used to enhance non-linear separability between vessel pixels and background, while batch normalization (BN) is utilized to stabilize training process under varying contrast conditions. The split features *g*_*h*_ and *g*_*w*_ represent orthogonal attention cues: *g*_*h*_ emphasizes horizontal vessel continuity (e.g., main arcades parallel to the retina’s equatorial plane), *g*_*w*_ highlights vertical branching patterns (e.g., perpendicular capillaries extending towards the macula). These directional attentions are obtained by element-wise multiplication which is designed to geometrically constrain vascular connectivity. Then they are subsequently fed into another two distinct conv layers to generate the attention maps. Lastly, the final re-weighted feature can be written as:

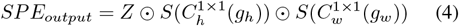

Here, *SPE*_*output*_ indicates the final output feature maps, while 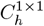 and 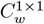 correspond to two distinct conv kernels of size 1×1. *S* represents the sigmoid gate, which is used to adjust feature responses and suppress disconnected pseudo-vessels while boosting true bifurcations according to anatomical plausibility.

#### 2) Transformer-derived Topology Interaction (TTI)

Although local trajectory continuity is ensured by the SPE module via axial-aware directional modeling, it does not explicitly address long-range topological dependencies between distant vessel branches, particularly at critical bifurcation junctions.

To overcome this limitation, we propose the Transformer-based Topology Interaction (TTI) module to explicitly model long-range spatial dependencies by incorporating vascular connectivity priors. Specifically, it constructs a pixel-wise affinity map by facilitating cross-region interactions among vascular segments, thereby allowing the model to: (a) improve bifurcation detection by amplifying structural associations between main vessels and their branches, and (b) suppress false discontinuities through performing long-range contextual reasoning along anatomically consistent paths. The transition from local morphology to global continuity enables the recovery of hemodynamically consistent vascular topologies. The designed topological interaction module, consisting of the following stages:

Firstly, two standard 3 × 3 convolutional layers are used to further encode *SPE*_*output*_ by fusing more richer local information. In order to capture orientation-specific contextual dependencies, the generated local feature 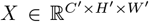 is then projected into three distinct representations—query *Q*, key *K*, and value *V* —throught the 1 × 3, 3 × 1, and 1 × 1 convolutional kernels, respectively. Here, these three convolutional kernels are explicitly used to model horizontal and vertical vascular trajectory continuity, aligning with the anisotropic structure of retinal structures. To enhance the inter-features relationships, they are then immediately flattened into a sequence of size *C*^*′*^ × *N* ^*′*^, where *N* ^*′*^ = *H*^*′*^ × *W* ^*′*^. A matrix multiplication is then performed to compute the affinity score between Q and K. The affinity score measures the probability of pairwise vascular connectivity, and the higher the value, the greater the likelihood of connection. Then, the softmax function is utilized to normalize the affinity score, resulting in the attention weights. Meanwhile, the value matric *V* is reshaped to 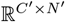 and is multiplied by the attention weights via a matrix multiplication, aiming to calculate the affinity attention maps. Finally, to avoid overemphasizing specific objects, we scale the affinity attention maps using a trainable parameter. These adjusted feature maps are subsequently combined with the original input *X* through a residual connection, resulting in improved generalization capacity for the model. The calculation method is defined by the following formula:

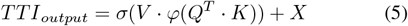

Here, *φ* denotes the softmax function for computing the attention weights, and · represents the matrix multiplication. The learnable scalar *σ* is used to balance global topology iniformation against local geometric accuracy, adapting to variations in vessel density across retinal regions.

#### 3) SSemantic Attention Amplification (SAA)

In retinal images, pathologies such as microaneurysms and hemorrhages, are characterized by localized intensity changes, whereas capillaries typically exhibit low-contrast patterns. Deep learning models inherently encode such variations into distinct channel features, enabling channel attention mechanisms to prioritize clinically salient regions [41], [42]. This selective focus enhances the model’s capability to effectively represent and emphasize functional semantics of inter-ichannel features. Thus, in this section, we further explore the inter-relationships among channel maps with the computed coordinate attention maps. Unlike the SE attention mechanism [23], which relies on scalar operations, our approach employs matrix multiplication to explicitly model the interactions between different channels. To quantify channel-wise semantic synergy, we firstly project the geometric-refined features 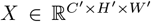 query *Q*^*′*^, Key *K*^*′*^ and Value *V* ^*′*^ via reshape operations. To obtain the channel affinity score Aff, we perform a multiplication between *Q*^*′*^ and the transposed *K*^*′*^. The higher affinity values indicate complementary channel pairs (e.g., microaneurysm channels and their surrounding capillary channels), whereas lower values suggest redundant or contradictory semantics. To amplify clinically critical channel interactions, we then apply contrast enhancement by subtracting each affinity value from the channel-wise maximum. T

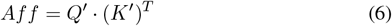

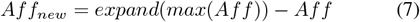

Where *expand* presents the *expand as* function, *max* presents the *max* function and 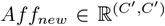 presents the resulting new affinity matrix. This operation suppresses dominant but less informative channel correlations (e.g., background-dominated channels), while highlighting subtle yet hemodynamic-indicative relationships. Similar to above section, we then normalize the calculated affinity matrix with softmax function *φ* and then it is multiped with *V* ^*′*^ by a linear matrix multiplication to produce the attention-weighted value matrix. In order to dynamically adjust the weights of the new attention feature map and the geometric-refined feature map *X*, we perform the following calculation formula:

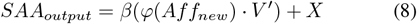

Where, *φ* denotes the softmax function, *SAA*_*output*_ represent the output of SAA module, and *β* represents the learnable parameter.

### C. Training Objectives

We employ binary cross-entropy (BCE) as the primary loss function for pixel-wise vessel segmentation:

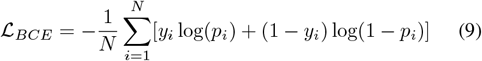

where *N* is the total number of pixels, *y*_*i*_ ∈ {0, 1} is the ground truth label, and *p*_*i*_ is the predicted vessel probability. This loss effectively handles the class imbalance inherent in retinal vessel segmentation.

### D. Architecture Variants

Although DFMS-Net attains high accuracy due to its integrated geometric-functional design, the distinct diagnostic tasks of real-world demand specialized architectures. To this end, two task-specific variants—Variant-1 and Variant-2—are proposed, which retain the fundamental architectural priors of DFMS-Net while being designed to tackle distinct clinical challenges in retinal vessel segmentation (see Appendix Fig A1).

Both variants are built upon a Double SPE Attention module within the bottleneck layer, which replaces the Dual-Field Hemodynamic Attention (SPE+TTI+SAA). Our design is motivated by ophthalmologists’ two-stage assessment: when assessing retinal images, clinicians do not rely on a single glance but conduct a thorough two-stage evaluation. Mirroring clinical practice, the model first identifies directionally coherent candidates from the input, then validate their topological continuity and connectivity by fusing contextual information (e.g., surrounding tissue contrast, branching patterns).

## IV. EXPERIMENTS

### A. Implementation Details and DATASETS

We evaluate our method on four public datasets with varying image resolutions and sizes, including DRIVE [43](40 images, 584×565), STARE [44](20 images), DCA1 [45] and CHUAC [46].

The proposed framework is implemented in PyTorch 2.0.1 on an NVIDIA A100 GPU. We employ the Adam optimizer to train the networks, with the learning rate and weight decay set to 0.0001 and 0.00001, respectively. Moreover, we utilize binary cross-entropy (BCE) as the loss function and adopt a cosine annealing learning rate scheduler. The learnable scaling parameters *σ* and *β* in both TTI and SAA modules are initialized to zero following the progressive learning principle, ensuring attention mechanisms start as identity mappings and gradually contribute as training progresses. This initialization strategy prevents attention collapse during early training while maintaining vessel connectivity preservation. Specifically, to conserve computational resources and augment data size, we extract image patches with a 48 × 48 sliding window using the unfold function. In this study, we evaluate the model’s segmentation performance by comparing the segmentation results to the ground truths using several widely used metrics, including the intersection over union (IoU), accuracy (Acc), sensitivity (Sen), specificity (Spe), F1 score (F1) and area under the curve (AUC). These criteria are defined as follows:

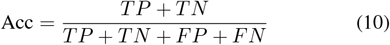

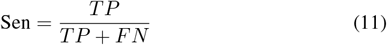

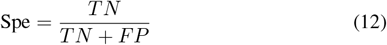

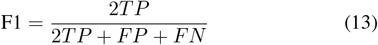

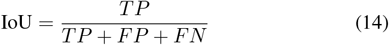

For medical segmentation results, pixels are divided into two categories: vascular pixels (those with blood flow) and non-vascular pixels (those without blood flow). The classification of each image produces four potential results: vascular pixels correctly identified as vessels (True Positive, TP), non-vascular pixels correctly identified as non-vessels (True Negative, TN), nonvascular pixels incorrectly predicted as vessels (False Positive, FP), vascular pixels incorrectly predicted as non-vessels (False Negative, FN).

### B. Comparative Experimental Studies

To evaluate the performance of our designed DFMS-Net, we conduct series experiments against state-of-the-art methods, including TransUNet [25], DE-DCGCN-EE [47], PLVS-Net, DPF-Net [28], MedSegDiff [27], GT-DLA-dsHFF [49] and DPL-GFT-EFA [28]. Also, we compare our framework with classic methods, such as U-Net [19], UNet++ [50] and Attention U-Net [13]. To demonstrate the advantages of our approach more effectively, we conduct a comprehensive evaluation incorporating both qualitative and quantitative analyses across four public benchmarks.

#### 1) Evaluation on the DRIVE Dataset

Our proposed DFMS-Net attains state-of-the-art performance on the DRIVE benchmark, showing a superior balance across key metrics. In TABLE I, our framework ranks first among all methods—accuracy (ACC) of 97.08%, specificity (Spe) of 98.52%, AUC of 98.89%, and IoU of 71.12%, exceling in suppressing false positives while keeping vascular continuity. Although GT-DLA-dsHFF [49] has a slightly higher sensitivity (Sen: 83.77% vs. 82.37%), the spatial redundancy introduced by hierarchical attention fusion, leads to a drop in Spe and IoU. The problem is solved by DFMS-Net through the special designed hemodynamic attention mechanisms, which incorporates the SPE module to ensure vessel trajectory smoothness and decrease fragmentation in low-contrast regions (Spe↑ 0.25% vs. GT-DLA-dsHFF). To solve the forking ambiguity problem, the designed TTI leverages adaptive affinity learning, obtaining the highest IoU of 71.12% and AUC of 98.89%. It should be highlighted that DFMS-Net surpasses the transformer-based TransUNet [25] by 0.82% and the diffusion-driven MedSegDiff [27] by 10.26% in IoU, verifying the promissing robustness to architecture-bias. While DPL-GFT-EFA [28] shows a slightly higher F1 score than DFMS-Net, its graph-based fusion has problems with capillary boundary stability (AUC: 98.69% vs. 98.89%). Meanwhile, the designed Variant-1 reaches almost the same level of performance with minimal reduction in parameters (IoU: 71.10%, Spe: 98.46%), by prioritizing symmetric-driven coherence. Through boundary-centric SPD [52], Variant-2 is improved by 0.93% in Sen compared to DFMS-Net, while keeping a robust IoU (71.06%). These outcomes validate DFMS-Net’s effectiveness in geometric precision and hemodynamics-related correlation.

**TABLE I.** Comparison with the state-of-the-art methods on DRIVE and STARE. The best results are in bold.

| Methods | DRIVE |  |  |  |  |  | STARE |  |  |  |  |  |
| --- | --- | --- | --- | --- | --- | --- | --- | --- | --- | --- | --- | --- |
|  | ACC (%) | Sen (%) | Spe (%) | AUC (%) | F1 (%) | IoU (%) | ACC (%) | Sen (%) | Spe (%) | AUC (%) | F1 (%) | IoU (%) |
| U-Net [19] | 96.78 | 80.57 | 98.33 | 98.25 | 81.41 | 68.64 | 97.30 | 78.50 | 98.84 | 98.75 | 81.18 | 68.56 |
| UNet++ [50] | 96.79 | 78.91 | 98.50 | 98.25 | 81.14 | 68.27 | 97.34 | 79.09 | 98.83 | 98.84 | 81.50 | 69.02 |
| Attention U-Net [13] | 96.62 | 79.06 | 98.31 | 97.74 | 80.39 | 67.21 | 97.30 | 78.04 | 98.87 | 98.78 | 81.06 | 68.39 |
| TransUNet [25] | 96.90 | 82.64 | 98.27 | 98.38 | 82.35 | 70.30 | 97.48 | 83.32 | 98.62 | 98.68 | 83.25 | 71.45 |
| DE-DCGCN-EE [47] | 97.05 | 83.59 | 98.26 | 98.66 | 82.88 | 70.77 | 97.51 | 84.05 | 98.61 | 98.99 | 83.63 | 71.87 |
| PLVS-Net [48] | 96.79 | 81.95 | 98.19 | 98.32 | 81.56 | 68.89 | 97.28 | 83.08 | 98.43 | 98.71 | 82.20 | 69.87 |
| DPF-Net [51] | 96.96 | 83.05 | 98.30 | 98.53 | 82.70 | 70.54 | 97.44 | 83.60 | 98.56 | 98.76 | 83.04 | 71.17 |
| MedSegDiff [27] | 95.61 | 78.00 | 97.31 | 94.32 | 75.65 | 60.86 | 91.22 | 70.56 | 92.83 | 88.42 | 66.43 | 50.80 |
| GT-DLA-dsHFF [49] | 97.00 | **83.77** | 98.27 | 98.66 | 82.99 | 70.96 | 97.60 | 84.88 | 98.65 | 99.13 | 84.32 | 72.89 |
| DPL-GFT-EFA [28] | 97.05 | 83.61 | 98.34 | 98.69 | **83.22** | 71.03 | 97.70 | 85.31 | 98.69 | 99.17 | 84.74 | 73.64 |
| Baseline | 97.05 | 83.13 | 98.40 | 98.87 | 83.07 | 71.06 | 97.65 | 87.39 | 98.67 | 99.42 | 86.75 | 76.70 |
| Variant-1 | 97.06 | 82.77 | 98.46 | 98.88 | 83.09 | 71.10 | 97.68 | 80.61 | 98.85 | 99.43 | 86.73 | 76.66 |
| Variant-2 | 97.04 | 83.30 | 98.38 | 98.87 | 83.06 | 71.06 | 97.71 | 85.55 | **98.92** | 99.44 | 86.83 | 76.80 |
| DFMS-Net | **97.08** | 82.37 | **98.52** | **98.89** | 83.11 | **71.12** | **97.72** | **88.01** | 98.68 | **99.47** | **87.18** | **77.35** |

#### 2) Evaluation on the STARE Dataset

On STARE, DFMS-Net surpasses current methods with a 0.62% higher sensitivity (88.01%) and a 0.65% higher IoU (77.35%), achieving a new top-level performance, with specificity (98.68%) and AUC (99.47%), which proves its diagnostic reliability. This breakthrough illustrates the framework’s ability in solving two lasting challenges in retinal analysis challenges: (a) the detection of low-contrast vessels and (b) the maintenance of peripheral vascular continuity. Although DPL-GFT-EFA [28] shows near-perfect discrimination (AUC 99.17%), its probability-based fusion strategy cannot capture the inherent anisotropic geometric prior in retinal vessels, leading to fragmented output in the macular region (F1: 84.74%). GT-DLA- dsHFF [49] achieves a high Sen score of 84.88%. However, its hierarchical attention blocks over-smooth fine vessels, leading to a 4.46% lower IoU than DFMS-Net. TransUNet [25] shows worse performance in Spe (98.62% vs. 98.68%) and F1 (83.25% vs. 87.18%) due to the lack of directional priors. In comparison, Variant-1 effectively balances IoU and Spe, while Variant-2 shows promising performance in boundary-critical situations, realizing 85.55% Sen (↑1.5% over DE-DCGCN-EE) using SPD-Conv’s spatial fidelity.

#### 3) Evaluation on the DCA1 and CHUAC Dataset

In coronary angiography, the proposed DFMS-Net shows significant break-through in consistency, which is evidenced by its remarkable performance on DCA1 and CHUAC benchmarks in Table II. The dual-field advantage verifies its practical ability to tackle two significant challenges in medical image segmentation: the detection of artifacts vessels and the preservation of continuity in low-signal pixel points. The proposed DFMS-Net reaches higher Spe on both datasets: 98.91% for DCA1 and 99.60% for CHUAC. Moreover, DFMS-Net demonstrates superior performance by focusing on the retention of vascular topology and enhancement of critical-vessel intensity, which is enabled by dynamic adjustment of the weights of geometric and branched structural features, thereby achieving a high AUC of 99.19%. It effectively suppresses noise interference with the phase-aware attention mechanism, obtaining a Sen of 82.12% on CHUAC, compared to UNet++’s 66.87%.

**TABLE II.**
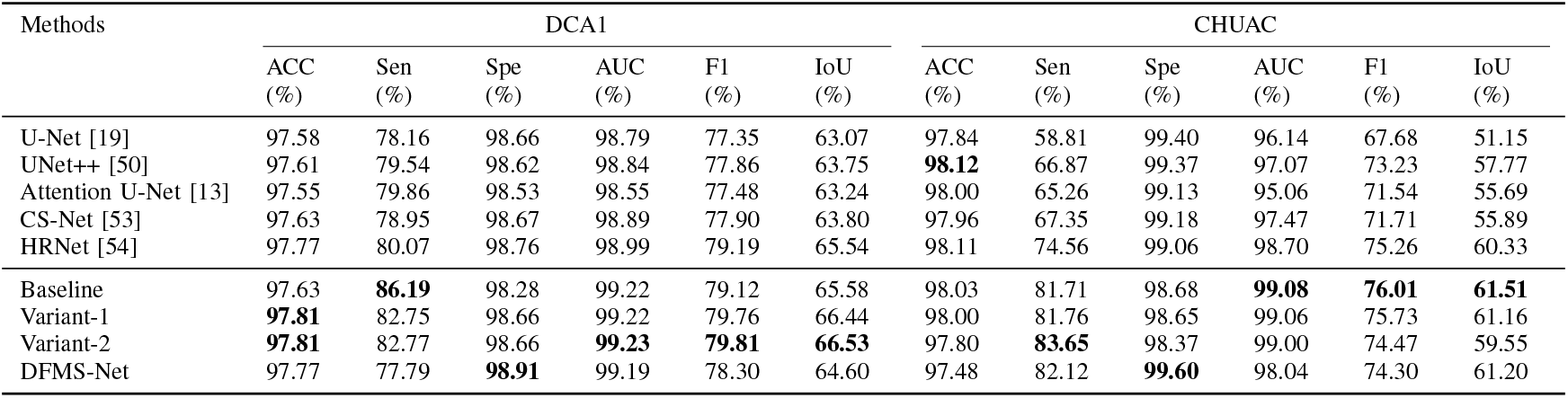
Comparison with the state-of-the-art methods on DCA1 AND CHUAC. The best results are in bold.

Variant-1 maintains a high ACC of 97.81% on DCA1, an increase of 0.18% compared to the baseline. Variant-2 achieves state-of-the-art IoU of 66.53% via the SPD-Conv’s non-destructive downsampling on DCA1, and obtains the highest Sen of 83.65% by preserving capillary endpoints in low-SNR regions on CHUAC, demonstrating strong advantage of recovering blurred boundaries and handling scenarios with significant vessel size variations.

#### 4) Visualization Segmentation Results

As visualized in Fig.2, the segmentation results show a clear demonstration of the dual-field synergy of DFMS-Net across retinal and coronary domains. Due to space limitations, we focus on comparing the baseline network with our proposed framework. In the retinal microvasculature (DRIVE and STARE), the baseline fragments delicate capillary networks—particularly in low-contrast periphery—as shown by discontinuous branching patterns (green arrows, rows 1 and 4), arising from its inability to model anatomical continuity under hemodynamic variations. DFMS-Net addresses this through its Spatial Pathway Extractor (SPE), which enforces trajectory-aware attention to bridge gaps between junctions while suppressing non-vessel responses (blue circle, row 1).

**Fig. 2.**
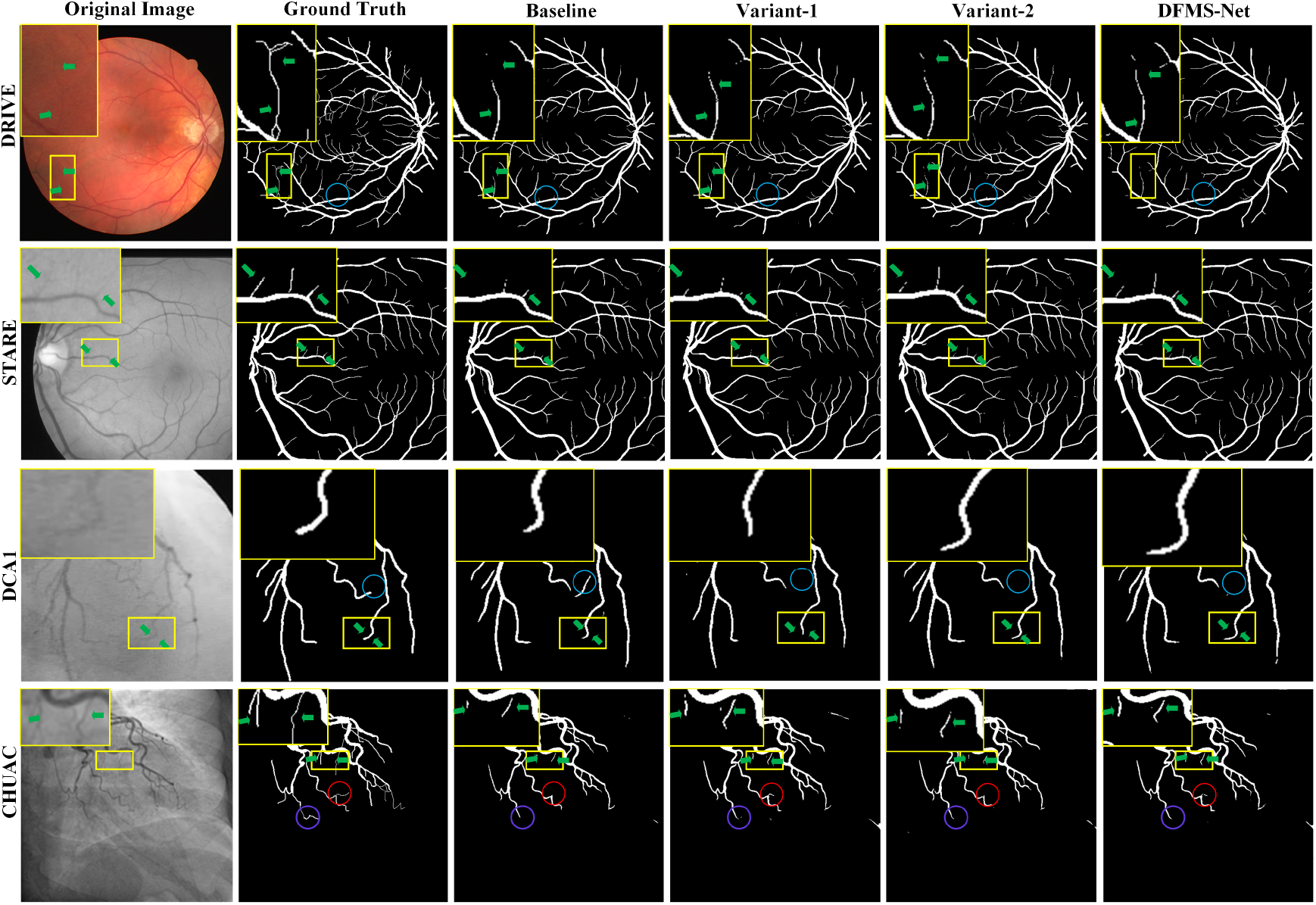
Segmentation comparison on four datasets. Yellow boxes show improved regions with zoomed views. Green arrows: capillary fragmentation in baseline. Blue circles: false positives suppressed by DFMS-Net (via SPE in retina, structural separation in coronary). Red circles: better branch recovery via SAA. Purple circles: Variant-2’s advantage in low-SNR areas using SPD-Conv. The results validate the dual-field design and variant-specific optimizations.

In the coronary domain (DCAI and CHUAC), the Semantic Attention Amplification (SAA) module enhances true capillaries over background textures, resulting in cleaner branch delineation (red circle, row 4). The framework effectively balances boundary sharpness and noise suppression, avoiding misclassification of pseudo-vessels (blue circle, row 3)—a critical failure mode of the baseline under static feature aggregation.

The variants further illustrate targeted improvements: Variant-1 preserves complex coronary structures with higher efficiency via SPE; Variant-2 excels in low-SNR sequences thanks to SPD-Conv, as evidenced by improved continuity in challenging regions (purple circle, green arrow, row 4).

### C. Ablation Study

To rigorously dissect the contributions of DFMS-Net’s dual-field paradigm, we perform systematic ablation studies on DRIVE and STARE datasets, which represent divergent vascular phenotypes (retinal microvasculature and complex bifurcations). We systematically disable individual modules (SPE/TTI/SAA), with configuration setting detailed in Table III. Table IV presents comprehensive quantitative metrics, while Fig. A3 in Appendix C anatomically dissect performance gains through color-coded segmentation results.

**TABLE III.** Abbreviations used in ablation studies.

| Abbreviation | Explanation |
| --- | --- |
| / | Parallel connection |
| SPD_down | Replacing strided convolution with non-strided convolution |
| Down_SPE | SPE module concatenated to standard down-sampling |
| Out_weight | Weighted sum of outputs |
| SPD_down_SPE | SPE module concatenated to SPD_down operation |
| Out_SPE | SPE module concatenated to output layer |

**TABLE IV.**
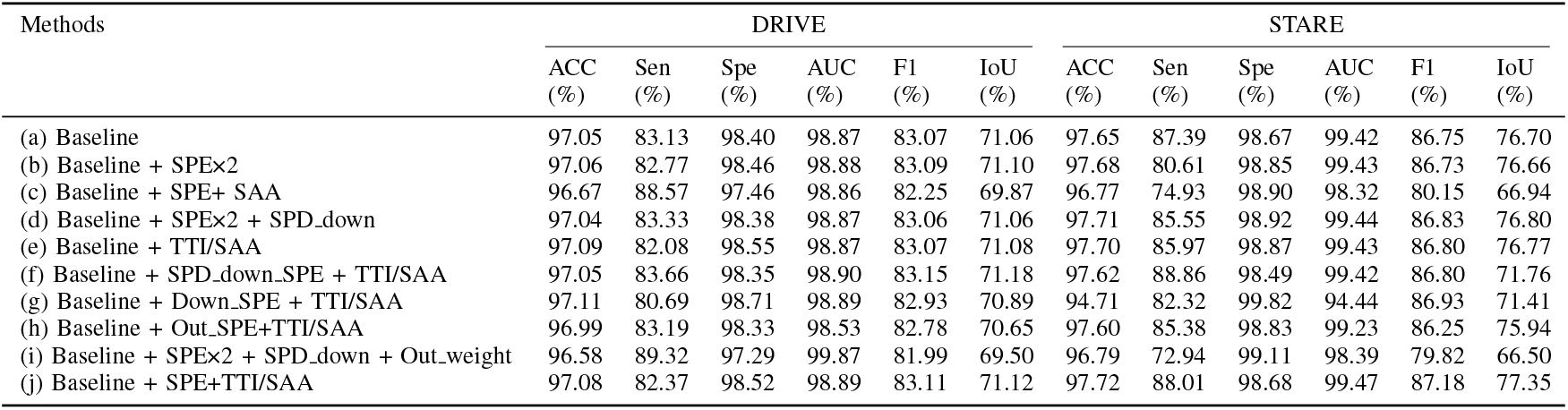
ABLATION EXPERIMENTS ON THE DRIVE AND STARE DATASETS.

| Methods | DRIVE |  |  |  |  |  | STARE |  |  |  |  |  |
| --- | --- | --- | --- | --- | --- | --- | --- | --- | --- | --- | --- | --- |
|  | ACC (%) | Sen (%) | Spe (%) | AUC (%) | F1 (%) | IoU (%) | ACC (%) | Sen (%) | Spe (%) | AUC (%) | F1 (%) | IoU (%) |
| (a) Baseline | 97.05 | 83.13 | 98.40 | 98.87 | 83.07 | 71.06 | 97.65 | 87.39 | 98.67 | 99.42 | 86.75 | 76.70 |
| (b) Baseline + SPE×2 | 97.06 | 82.77 | 98.46 | 98.88 | 83.09 | 71.10 | 97.68 | 80.61 | 98.85 | 99.43 | 86.73 | 76.66 |
| (c) Baseline + SPE+ SAA | 96.67 | 88.57 | 97.46 | 98.86 | 82.25 | 69.87 | 96.77 | 74.93 | 98.90 | 98.32 | 80.15 | 66.94 |
| (d) Baseline + SPE×2 + SPD_down | 97.04 | 83.33 | 98.38 | 98.87 | 83.06 | 71.06 | 97.71 | 85.55 | 98.92 | 99.44 | 86.83 | 76.80 |
| (e) Baseline + TTI/SAA | 97.09 | 82.08 | 98.55 | 98.87 | 83.07 | 71.08 | 97.70 | 85.97 | 98.87 | 99.43 | 86.80 | 76.77 |
| (f) Baseline + SPD_down.SPE + TTI/SAA | 97.05 | 83.66 | 98.35 | 98.90 | 83.15 | 71.18 | 97.62 | 88.86 | 98.49 | 99.42 | 86.80 | 71.76 |
| (g) Baseline + Down.SPE + TTI/SAA | 97.11 | 80.69 | 98.71 | 98.89 | 82.93 | 70.89 | 94.71 | 82.32 | 99.82 | 94.44 | 86.93 | 71.41 |
| (h) Baseline + Out.SPE+TTI/SAA | 96.99 | 83.19 | 98.33 | 98.53 | 82.78 | 70.65 | 97.60 | 85.38 | 98.83 | 99.23 | 86.25 | 75.94 |
| (i) Baseline + SPE×2 + SPD_down + Out_weight | 96.58 | 89.32 | 97.29 | 99.87 | 81.99 | 69.50 | 96.79 | 72.94 | 99.11 | 98.39 | 79.82 | 66.50 |
| (j) Baseline + SPE+TTI/SAA | 97.08 | 82.37 | 98.52 | 98.89 | 83.11 | 71.12 | 97.72 | 88.01 | 98.68 | 99.47 | 87.18 | 77.35 |

#### 1) Network Configurations

To deconstruct the contributions of DFMS-Net’s dual-field components, we engineer ten progressively enriched architectures atop the FR-UNet baseline, by systematically integrating modules from the geometric (SPE, TTI) and functional (SAA) fields. Configurations are codified in Table III, with key variants categorized as: (a) Baseline: FR-Unet network. (b) Baseline + SPE×2 (Variant-1): The architecture includes two SPE modules based on baseline network. (c) Baseline + SPE+ SAA: It means that the architecture is composed of one SPE module and one SAA module based on baseline network. (d) Baseline + SPE×2 + SPD down (Variant-2): It means that the architecture is composed of two SPE modules and one SPD down module based on baseline network. (e) Baseline + TTI/SAA: It represents that the TTI and the SAA modules are paralleled designed based on the baseline network. (f) Baseline + SPD down SPE + TTI/SAA: It represents that the network consists of SPD down, SPE, TTI and SAA components based on the baseline network. (g) Baseline + Down SPE + TTI/SAA: It presents that the network consists of Down SPE, TTI and SAA components based on the baseline network. (h) Baseline + Out SPE+ TTI/SAA: It represents that the network consists of Out SPE, TTI and SAA components based on the baseline network. (i) Baseline + SPE×2 + SPD down + Out weight: It represents that the network consists of two SPE, one SPD down and one Out weight modules based on the baseline network. (j) Baseline + SPE + TTI / SAA (DFMS-Net.): It represents that the network consists of SPE, TTI and SAA modules based on the baseline network.

#### 2) Effect of the SPE module

The SPE module integrates geometric accuracy and hemodynamics-relatedness through orientation-aware coordinate attention, as demonstrated in TABLE IV. On the DRIVE dataset, incorporating SPE into the network (j) improves sensitivity by 0.2% and IoU by 0.04% over network (e), which uses only TTI and SAA modules. Although these gains may appear modest in absolute terms, they are statistically significant given the high complexity and low contrast of retinal microvasculature. As visualized in the yellow box in Fig. A3 (Appendix C), network (j) substantially reduces false negatives (misclassified red pixels) and recovers fine capillaries that are typically lost due to background noise—validating the role of SPE in enhancing structural continuity and noise resilience.

The results on STARE dataset further demonstrate the capability of SPE. Specifically, network (j) obtains an improvement of 2.04% in Sen over network (e), validating its superior performance in identifying thin-vessels branches. Notably, SPE addresses a key limitation of TTI and SAA: while these modules excel at global context aggregation, they fail to preserve hemodynamic-geometric relationships critical for fine-scale vascular continuity.Furthermore, ablation studies reveal that integrating SPE in the encoding stage yields greater performance gains than in the decoding stage, indicating its role in early structural encoding. This is evidenced by an F1 score of 83.11% on DRIVE and 87.18% on STARE when SPE is applied in the encoder (network (h) vs. (j)).

#### 3) Effect of the TTI module

On the DRIVE dataset, the TTI module reduces edge ambiguity in thick vessels by modeling spatial connectivity priors, as shown in the blue box in Fig. A3 (Appendix C, network (j) vs. (c)). This enables clearer separation of adjacent vessel boundaries (green box) and yields a 1.25% improvement in IoU (71.12% vs. 69.87%, Table IV). Although sensitivity increases only marginally (88.57% vs. 82.37%), network (c) struggles to balance geometric precision with diagnostic fidelity, often over-smoothing fine structures or misclassifying ambiguous edges.

Notably, on the STARE dataset, TTI boosts sensitivity by +13.08% over network (c) (SPE+SAA), demonstrating its ability to recover missed vessels through sequential topological reasoning. This improvement is primarily arises from the successful reconstruction of broken vascular bifurcations that are typically severed by noise or low contrast in conventional methods. This improvement is further evidenced by the substantial 10.41% absolute gain in IoU—from 66.94% (network c) to 77.35% (network j), reducing false gaps and fragmented outputs that could mislead diagnosis.

#### 4) Effect of the SPD down module

Experiments across topological configurations (b)-(d) and (g)-(f) in Table IV, reveals the SPD down’s critical advantages. The IoU metric demonstrates its stability on DRIVE (71.10% → 71.06%), comparing network (b) with (d) (red box, Fig. A3 in Appendix C). Yet it shows an increase on STARE (76.66% → 76.80%), indicating improved adaptability to complex vascular structures. Meanwhile, the designed module illustrates its enhanced detection of capillaries without increasing false positives on STARE (Spe: 98.85% → 98.92%, Sen: 80.61%→85.55%). Similarly, network (f) demonstrates a substantial improvement in Sen over (g), showcasing superior detection capability for low-contrast lesions. As illustrated in Fig. A3 in Appendix C, the SPD-integrated method (network (f)) notably improves small vessel detection accuracy. Concurrently, it effectively reduces the misclassification of non-vessel pixels(red and blue boxes, row 2).

#### 5) Effect of the output strategy module

To address the inherent class imbalance in retinal vasculature segmentation (vessel-to-background pixel), we rigorously evaluate two hierarchical fusion strategies in Table IV: (a) Deep Supervision with weighted summation (network (i)), which aggregates features from the last five layers using learnable weights, and (b) Average fusion (network (d)), where the multi-outputs are combined through element-wise averaging. On DRIVE, weighted summation approach (network (i) achieves higher Sen and AUC compared to network (d), but sacrifices ACC(↓0.46%), Spe (↓1.09%), F1(↓1.07%) and IoU(↓1.56%). This trade-off suggests potential overfitting to vessel pixels in class-imbalanced scenarios. On STARE, the average fusion strategy dominates across all metrics (ACC: ↑0.92%, Sen: ↑12.61, AUC: ↑1.05%, F1: ↑7.01%, IoU: ↑10.30%), demonstrating better generalization on complex vascular networks.

The limited capability of deep supervision can be attributed to inherent gradient inconsistency. During backpropagation, each layer parameter has the effect of increasing the slight feature differences in low-contrast regions. Conversely, average fusion method keeps angiographic coherence, which is prominently illustrated by the increased intensity in the same spatial area Fig. A3 in Appendix C (network (d) vs. (i)).

### D. Topological Performance Analysis

Vascular connectivity preservation is essential for accurate hemodynamic analysis and disease diagnosis, as minor topological errors can significantly impact subsequent clinical quantification. To comprehensively evaluate the topological preservation capabilities of our dual-field framework, we conducted extensive comparisons with state-of-the-art methods using established topology-specific metrics [16], [18]: clDice for connectivity preservation, Betti 0 error (*β*_0_) measuring connected component differences, and Betti 1 error (*β*_1_) measuring topological hole differences in vascular networks.

#### 1) Benchmarking Topological Performance

Fig. 3 evaluates the topological performance of DFMS-Net and its ablated variants. The full model reduces *β*_1_ separation errors from 38.1% to 15.0%, a 60.6% relative reduction, indicating enhanced preservation of fine-scale vessel continuity—critical for detecting early microvascular abnormalities such as capillary non-perfusion and microaneurysms in diabetic retinopathy (DR). *β*_0_ connection errors decrease from 79.7% to 67.8% (11.9-point improvement), and the clDice score reaches 81.37%, outperforming both the baseline (81.32%) and the original clDice method (80.95%) [16]. Compared to TopoMortar (*β*_0_ = 61.04 ± 8.89) [18], our method achieves comparable *β*_0_ performance with significantly lower *β*_1_ errors.

**Fig. 3.**
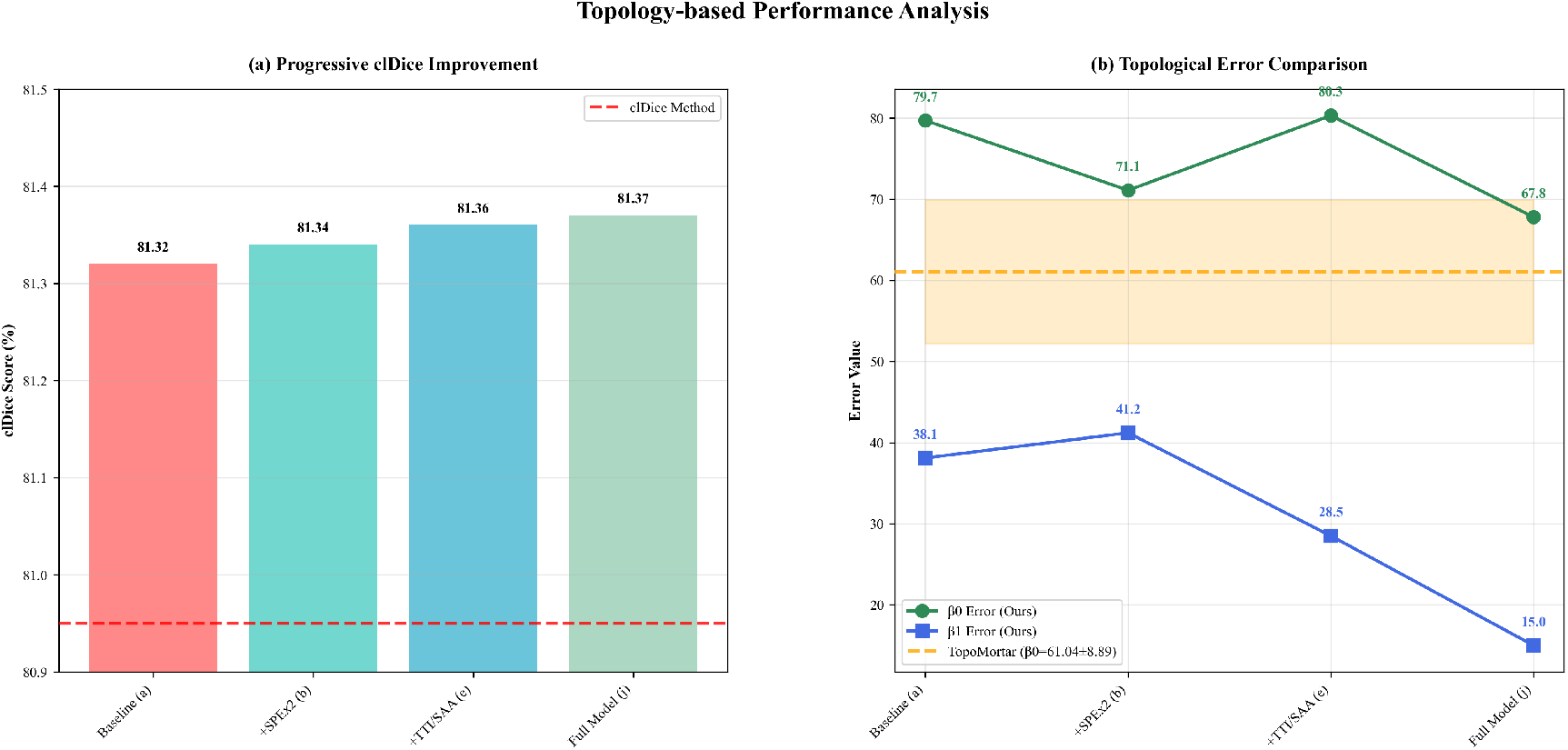
Benchmarking Topological Performance. (a) clDice scores with progressive addition of components: baseline (a), +SPE ***×*** (b), +TT/SAA (e), and full model (j) (DFMS-Net), compared with the clDice method. (b) ***β***_**0**_ (green) and ***β***_**1**_ (blue) topological errors across different models.

##### Stage 1 – Adding SPE

Incorporating the Spatial Pathway Extractor (SPE) reduces *β*_0_ errors from 79.7% to 71.1%, reflecting improved geometric continuity. However, *β*_1_ errors increase slightly to 41.2%, suggesting potential over-smoothing in complex regions that may obscure subtle pathological changes. The clDice score increases marginally to 81.34%.

##### Stage 2 – Adding TTI and SAA

The integration of TTI and SAA modules reduces *β*_1_ errors to 28.5% with stable *β*_0_ performance (80.3%),, indicating effective correction of local topological inconsistencies. Notably, SAA enhances semantic coherence across vessel scales, which is essential for distinguishing pathological vessel remodeling from normal branching patterns—a key challenge in early DR diagnosis.

##### Stage 3 – Full Integration (DFMS-Net)

The integration of SPE, TTA and SAA enables DFMS-Net to yields the minimal *β*_0_ (67.8%) and *β*_1_ (15.0%) errors the maximal clDice (81.37%). This three-part architecture reflects the step-by-step reasoning of ophthalmologists—first identifying vessel paths (SPE), then validating their continuity (TTI), and finally interpreting them within local tissue context (SAA)—leading to robust and comprehensive vascular structure preservation. From a clinical standpoint, the accurate preservation of vascular connectivity supports reliable analysis of microvascular health—thereby forming the foundation for pre-symptomatic detection of diabetic retinopathy.

#### 2) Mechanistic Analysis of Module Contributions

To clarify the functional roles of each component in our framework, we conduct a mechanistic analysis focusing on how SPE, TTI, and SAA address distinct topological challenges. The SPE module strengthens geometric continuity by selectively amplifying responses along dominant vessel directions. The TTI module identifies and rectifies local topological errors such as broken branches or false connections via global context modeling. Meanwhile, vessel semantics of different scales are preserved by SAA module through selective enhancement of features corresponding to fragile vessels, especially those with small diameters or low contrast. Collectively, Fig. 4 and 5 offer complementary quantitative and visual support of how the proposed modules collaboratively refine vascular topology by specialized mechanisms.

**Fig. 4.**
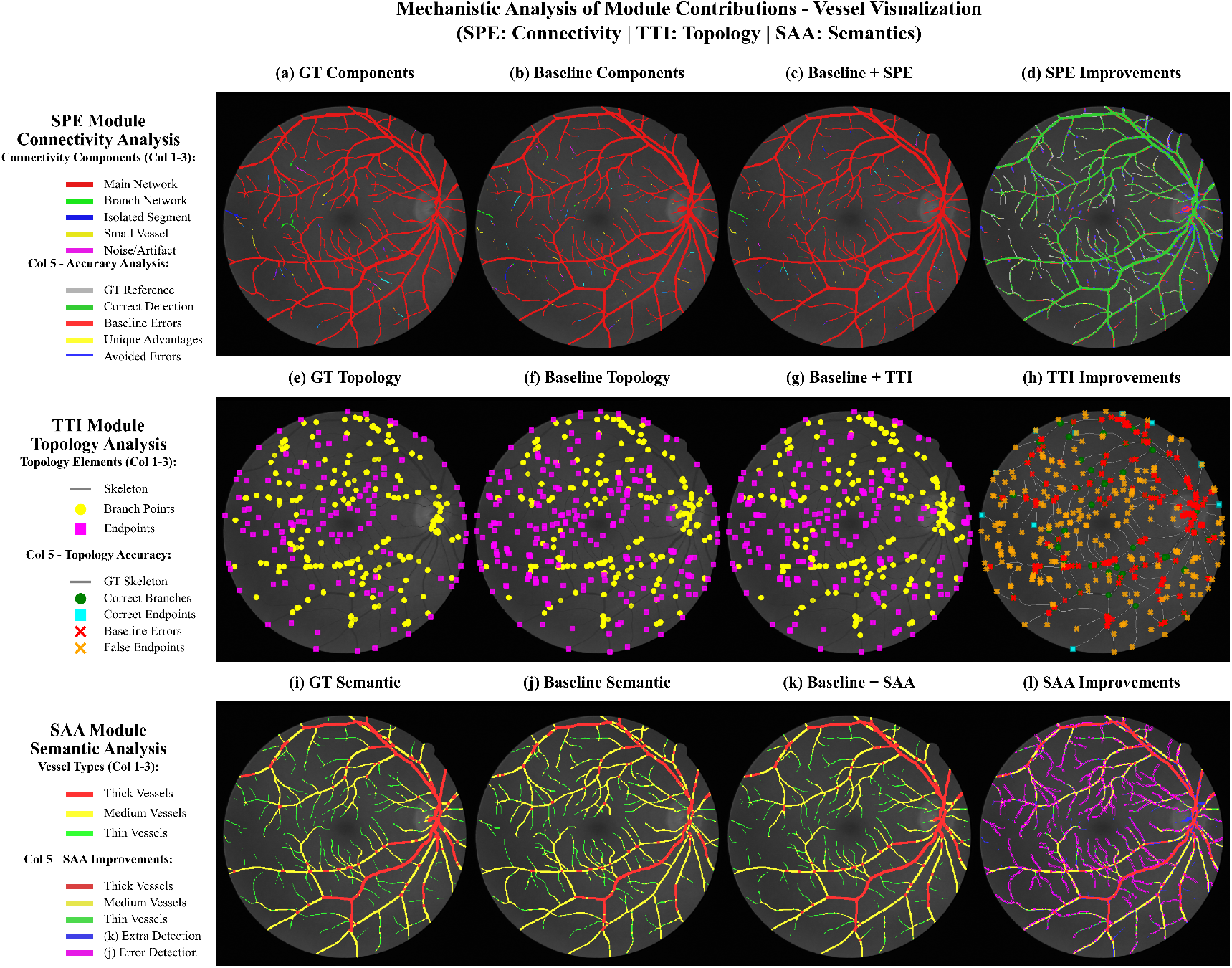
Analysis of Individual Module Contributions. Rows 1-3 show the outputs for the SPE, TTI, and SAA modules, respectively. Each row compares (Col 1) Ground Truth, (Col 2) Baseline limitations, (Col 3) Enhanced result, and (Col 4) Improvement map. The sequence SPE→TTI→SAA visually validates the quantitative gains in connectivity, topology, and semantics.

##### SPE Module - Connectivity Enhancement Analysis

As shown in Fig. 4, the baseline model produces isolated fragments predictions along vessel boundaries (blue), whereas the integration of SPE promotes path coherence by linking these disconnected components to the main or branch vessels (green extensions), thereby enhancing structural continuity. Moreover, quantitative results in Fig. 5(a) supports the visual improvement: the SPE module increases the connectivity quality score from 72.5% (baseline) to 87.2%, nearing the expert-level consistency (98.2%) essential for identifying early capillary dropout in DR. Meanwhile, boundary precision is increased from 68.3% to 85.6% and noise suppression is boosted from 75.2% to 89.4%, leading to a marked reduction in false positives and improving reliability in automated diabetic retinopathy screening. This shows that SPE not only refines vessel boundaries but also suppresses false detections. These statistical improvements offer quantitative validation for the visual reduction in vessel fragmentation and improved structural continuity seen in Fig. 4, and help account for the significant drop in *β*_0_ from 79.7% to 71.1%.

**Fig. 5.**
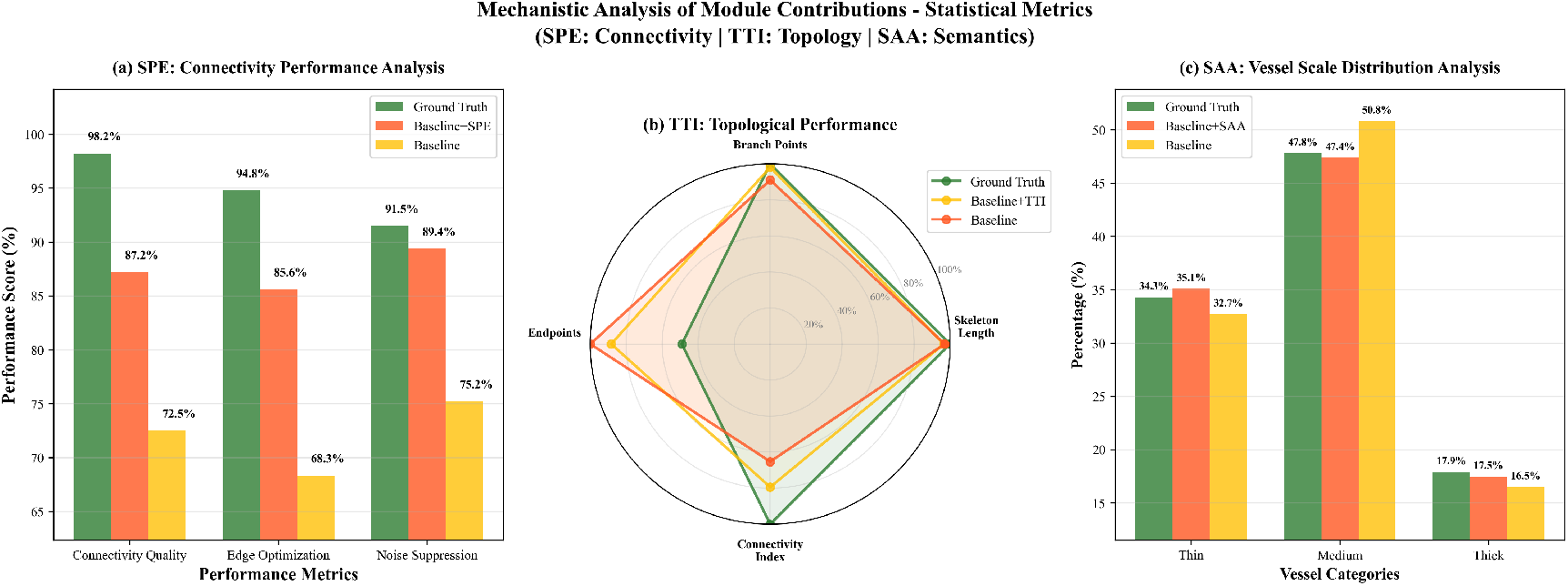
Quantitative performance validation of module-specific contributions. Statistical metrics align with Fig. 4: (a) SPE significantly boosts connectivity; (b) TTI improves topological structure; (c) SAA enhances multi-scale vessel detection. These results provide direct quantitative support for the module contributions.

Importantly, SPE’s role extends beyond simple “smoothing” or “filling”: it enables selective geometric completion through direction-aware feature propagation during the encoding stage. As evident in Fig. 4(d), green denotes correct reconnections and blue shows suppressed false connections—demonstrating its ability of topological discrimination.

##### TTI Module - Topological Structure Optimization

The TTI module is designed to identify and rectify structural errors-such as broken paths, spurious branches, or false end-points. Fig. 4(e–g) shows that the baseline model predicts excessive erroneous branch points (yellow circles) and endpoints (magenta squares), reflecting its limited capacity to preserve the structure of blood vessels. By modeling global contextual relationships, TTI resolves these topological errors-enabling the identification and correction of structural inconsistencies in the vascular network. Fig. 4(h) visualizes the baseline errors with TTI-corrected structures:red crosses indicate baseline errors, and orange crosses mark false endpoints, while green circles indicate recovered correct branches and cyan squares mark corrected endpoints after TTI refinement. Meanwhile, the radar chart in Fig. 5(b) demonstrates that Baseline+TTI exhibits superior performance, closely matching the ground truth results across multiple topological metrics-particularly in Branch Points, Endpoints, and Connectivity Index.

The improved performance indicates the strong ability of TTI to enhance anatomical plausibility and structural consistency of vascular topology. This correction enables more accurate localization of vessel branch points, which supports the early detection of retinal microangiopathy, where abnormal vascular bifurcations serve as key early biomarkers.

##### SAA Module - Semantic Detail Enhancement

Fig. 4(i)– (l) reveals how the SAA module improves semantic consistency across varying vessel scales and intensities. The GT Semantic labels (GT Semantic) are semantically classified into three classes: thick (red), medium (yellow) and thin (green), reflecting the multi-scale architecture of the retinal vascular vessels. Fig. 4(j) demonstrates significant under-segmentation of thin vessels, leading to extensive absence of green-labeled capillaries. Moreover, the baseline model shows significant semantic confusion in vessel scales where both thin and thick vessels are erroneously misclassified to the medium class (yellow), resulting in the excessive expansion of yellow areas. This semantic errors may lead to false-negative diagnoses or misinterpretation of disease severity. The incorporation of the SAA module (Fig. 4(k)) generates significant improvement in the detection and continuity of thin vessels, with restored capillary structures at the optic disc edge and periphery. The difference map (Fig. 4(l)) further corroborates this finding, which mark corrected vessel segments that were previously misclassified by the baseline.

Quantitative results are summarized in Fig. 5(c). The baseline model exhibits a strong bias toward medium-sized vessels (50.8%), while underestimating thin (32.7%) and thick (16.5%) vessels. The SAA-integrated model produces a more balanced vessel caliber distribution: 35.1% (thin), 47.4% (medium), and 17.5% (thick)—all in close agreement with the ground truth (34.3%, 47.8%, 17.9%). Therefore, the SAA module successfully preserves global semantic consistency by leveraging semantic-guided feature re-weighting, which is essential—for remaining anatomically and pathologically meaningful across the entire retinal field—and modeling the heterogeneous vascular pathology in diabetic retinopathy.

## V. DISCUSSIONS

Our introduced DFMS-Net framework demonstrates a novel integration of hemodynamic modeling with deep neural networks through the dual-field hemodynamic attention component—SPE and TTl for geometric-field modeling, while the SAA for functional-field optimization. By embedding topological information into the network, it achieves significant performance not only in conventional segmentation accuracy metrics (e.g., AUC, F1-score) but also in vascular topology metrics—such as clDice, *β*_0_ and *β*_1_—key biomarkers for staging and treatment planning—which are critical for reliable morphometric analysis in diabetic retinopathy, where capillary non-perfusion and microvascular dropout serve as early biomarkers.

Experimental results confirm these improvements across multiple datasets. Notably, on DRIVE, connectivity preservation improves to 81.37% in clDice (vs 80.95% in the baseline), with dramatic reductions in topological errors (*β*_0_: 67.8% vs 79.7%; *β*_1_: 15.0% vs 38.1%). The clinical value of the improved topological fidelity stems from its ability to preserve anatomical integrity, with key structural features such as connectivity, bifurcation patterns, and structural coherence. By correcting topological errors such as false endpoints, missing branches and broken segments (illustrated in Fig.4), our approach enables more precise computation of vascular indices such as connectivity quality and vessel categories—critical for monitoring diabetic retinopathy progression. Notably, these reductions in topological artifacts lead to more efficient and reproducible quantification of vascular biomarkers, providing a more trustworthy foundation for downstream morphometric analysis in clinical studies.

Despite promising performance, several limitations need careful consideration particularly for clinical practice. First, from the model design perspective, our framework lacks comprehensive cross-path information interaction—such as cross-attention or adaptive gating—between the encoder, decoder, and attention modules. This fusion strategy restricts bidirectional contextual enhancement and prevents the decoder from modulating the encoder-side features, constraining the effective use of complementary hierarchical representations. Second, the model does not include specialized modules to suppress non-vascular structures (e.g., exudates, hemorrhages, or artifacts), making it vulnerable to misclassifying non-vascular patterns as vessel pixels. Additionally, the current framework operates only on structural fundus images,lacking of functional modalities datasets. Therefore, the model demonstrates strong performance on the specific training domain (e.g., DRIVE), while shows performance degradation when applied to images from different domains with varying acquisition protocols, resolutions, or pathological distributions. The clinical interpretability of these metrics depends on their longitudinal stability. To demonstrate clinical effectiveness, it is necessary to conduct longitudinal studies to validate whether the proposed topological metrics correlate with clinically observed disease progression.

Moving forward, we aim to advance DFMS-Net by developing a unified framework for clinical deployment. we will focus on: (1) exploration of cross-attention interaction between structural and functional contextual information.generalization across various imaging protocols and pathological conditions through (2) domain adaptation techniques; (3) longitudinal evaluation of integrated structural and functional metrics to assess their sensitivity to disease progression and treatment response. These efforts aim to establish DFMS-Net as a clinically impactful tool, guiding early detection, staging, and individualized monitoring of microvascular disorders.

## VI. CONCLUSIONS

This work presents DFMS-Net, a novel framework integrating introduces a dual-field hemodynamic attention mechanism for retinal vessel segmentation, integrating geometric-field modeling and functional-field optimization. By embedding physiological information, our approach achieves superior segmentation accuracy and topological preservation. Beyond performance gains, our approach enables physiologically consistent predictions that support clinical interpretation. Future work will leverage cross-domain vascular mapping across various datasets to enhance generalization capabilities, enabling robust and consistent morphometric analysis in diabetic retinopathy, hypertensive retinopathy and retinal vein occlusion. By bridging domain gaps in imaging and pathology, this direction aims to transform DFMS-Net into a reliable system for automated biomarker extraction, supporting individualized patient management.

## VII. ACKNOWLEDGMENT

This work was supported by the Science and Technology Development Fund of Macao under Grant No. 0002/2024/RIA1, and by Fujian Province Young and Middleaged Teachers’ Educational Science Research Projects (Science and Technology Category) under Grant No. JAT232014, and by Research Projects of Putian University 2023036 and 2022047, and by Putian City Science and Technology Bureau under Grants 2022SZ3001ptxy03, and by Putian Science and Technology Plan Project 2023GJGZ003.

## APPENDIX

### A. Architecture Variants

We present detailed architectural descriptions of Variant-1 and Variant-2, which are derived from DFMS-Net but optimized for distinct clinical tasks in retinal vessel segmentation,as illustrated in Appendix Fig. A1. To better model the directionality-dependent structural degradation in retinal ischemia and glaucoma, Variant-1 is introduced through dual-stage refinement. Variant-2 is tailored for high-resolution representation of capillary dropout in early diabetic retinopathy, recovering detailed vessel information via SPD-Conv.

#### 1) Variant-1

Variant-1 replaces the full Dual-Field Hemodynamic Attention (SPE, TTI, and SAA)—with a simplified dual-stage SPE attention block within the bottleneck layer. The initial SPE module processes the feature map *Z* from the encoder stage and captures preliminary directional responses along horizontal and vertical, forming a foundation for subsequent trajectory refinement followed by two lightweight 3 × 3 convolution for local contextual aggregation. Subsequently, a second SPE module is employed to refine feature representation and reinforce trajectory consistency, emulating the multi-stage clinical reasoning process. This cascaded design improves vessel continuity, especially in radially organized regions such as the peripapillary area—critical for detecting structural disruptions in glaucoma.

#### 2) Variant-2

Variant-2 builds upon Variant-1 by replacing strided convolutions with non-stride convolution (SPD-Conv) in the encoder and decoder pathways, preserving fine spatial details in downsampling without compromising efficiency. Unlike traditional strided convolutions, which discard boundary voxels and reduce resolution through spatial compression, SPD-Conv employs a two-step strategy (Appendix Fig. A2) : first, a 1× 1 convolution is used to adjust channel dimensions without changing the spatial resolution; second, the feature map *Y* ∈ ℝ ^(*D,S,S*)^ is decomposed into four sub-features via stride-2 slicing:

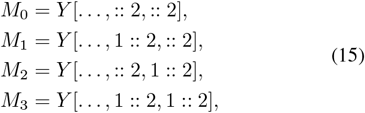

where each sub-feature captures spatially interleaved patches of the original map. These are then concatenated along the channel dimension to form a downsampled representation *Y*_out_ ∈ ℝ ^(4*D,S/*2,*S/*2)^, effectively halving the spatial resolution while quadrupling the channel dimension. As a result, SPD-Conv enables finer-grained feature abstraction without destructive spatial compression, allowing Variant-2 to recover faint vessels and capillary networks even in low-contrast images. Combined with the same Double SPE Attention module as in Variant-1, this design ensures both macro-scale directional consistency and micro-scale boundary integrity, making it ideal for high-resolution microvascular analysis such as early diabetic retinopathy grading and capillary density mapping.

**Fig. A1.**
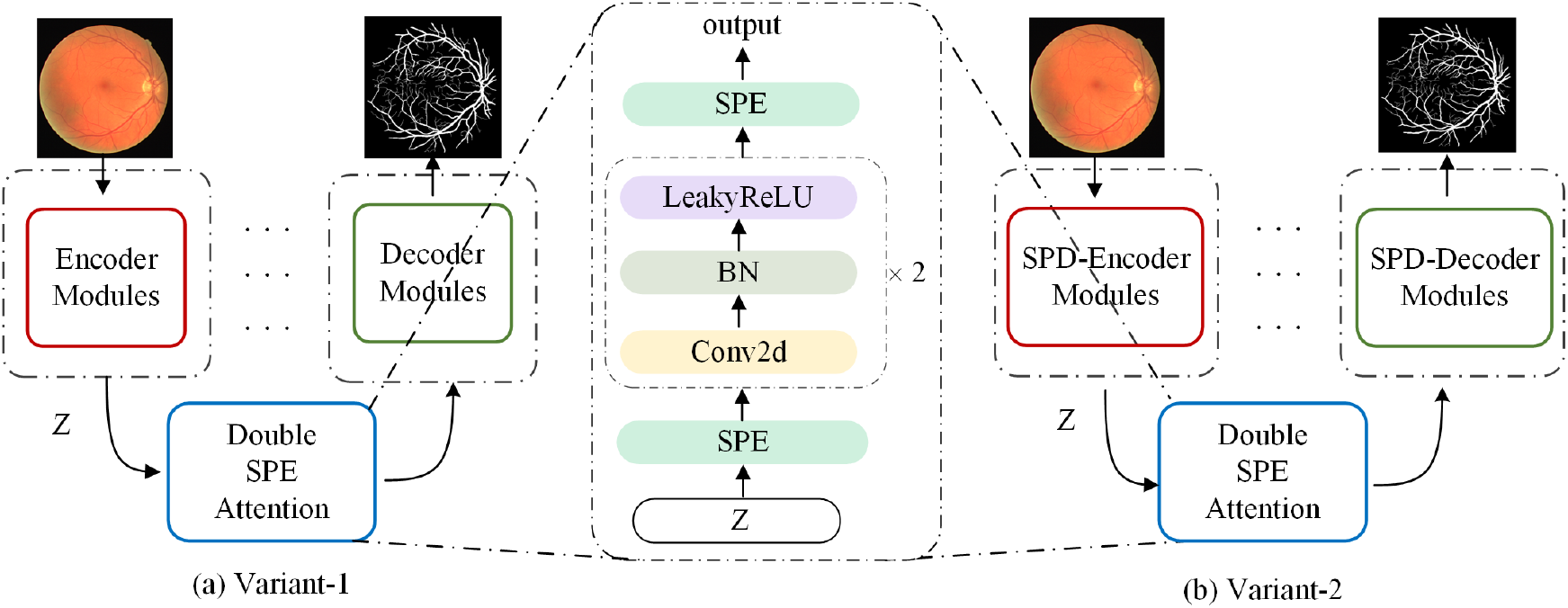
Architecture comparison of Variant-1 and Variant-2. (a) Variant-1 uses standard encoder-decoder blocks with a Double SPE Attention module in the bottleneck. (b) Variant-2 replaces them with SPD-Encoder and SPD-Decoder to preserve fine-scale details via sub-pixel downsampling. Both share the same attention mechanism for directional coherence.

**Fig. A2.**
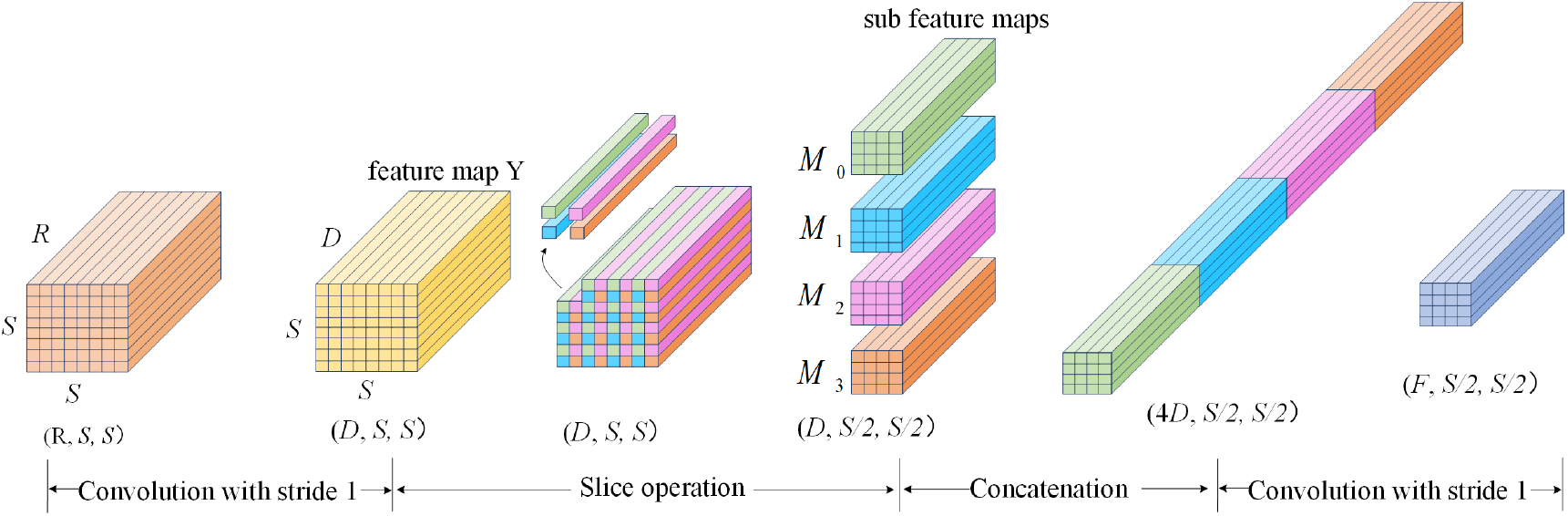
SPD-down module. The newly designed SPD downsampling operation is based on slicing strategy. We firstly use the convolution layer with stride 1 to change channel numbers, then the slice operation is utilized to generate four sub feature maps. Finally, the convolution layer is reused to decrease channel numbers.

### B. Visualized colored segmentation results

As a supplementary to the quantitative results in the main text, Fig. A3 (in Appendix) provides the visual segmentation results of the ablation study on the DRIVE dataset. The performance of different network variants, labeled from (a) to (j), is compared. In these visualizations, fluorescent blue indicates correctly segmented vessel pixels, while red and yellow denote incorrectly segmented vessels and background pixels, respectively.

**TABLE A1.** Computational Efficiency Analysis of Ablation Study Components.

| Model Configuration | Params (M) | FLOPs (G) |
| --- | --- | --- |
| Baseline | 5.72 | 2.07 |
| Variant-1 | 5.73 | 2.07 |
| Variant-2 | 5.84 | 2.07 |
| DFMS-Net | 5.76 | 2.08 |

**Fig. A3.**
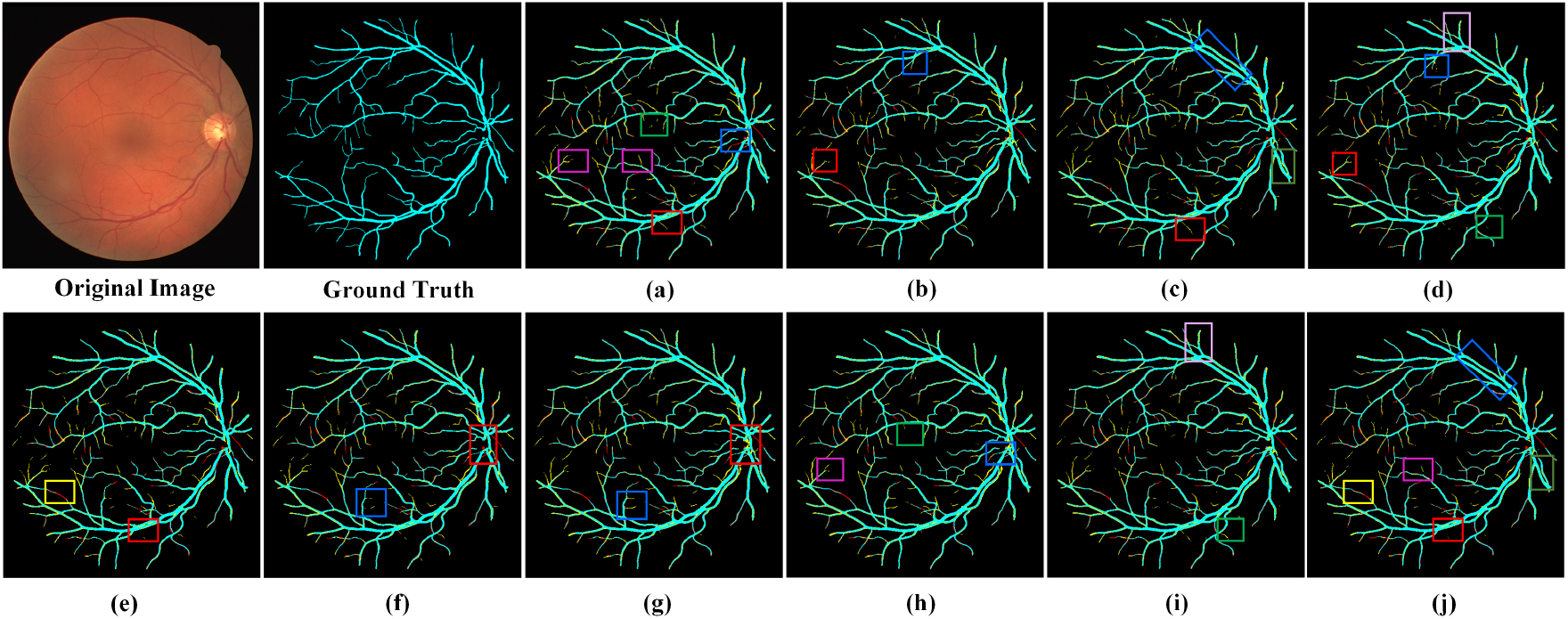
Visualized colored segmentation results of ablation study on DRIVE dataset of network (a)-(j). The fluorescent blue pixels present the correctly segmented vessel pixels, the red and yellow pixels present the incorrectly vessel pixels and background pixels, respectively.

### C. Model Computational Efficiency

The proposed architectures (Variant-1, Variant-2, and DFMS-Net) are designed to enhance segmentation accuracy with minimal increase in model size or computational burden. As shown in Table A1 (in Appendix), all variants exhibit minimal parameter growth (5.73–5.84million)—slightly above the baseline (5.72 million). Despite the integration of the dual-field hemodynamic attention to timprove vessel continuity and capillary recovery, yet their inference FLOPs remain at 2.07G and 2.08G, respectively—comparable to the baseline. Even Variant-2 which supports high-resolution analysis of microvascular, introduces just 0.11M extra parameters and maintains the same inference FLOP count as the baseline. These results demonstrate that the performance gains in Section IV are attributable to intelligent architecture design rather than increased model capacity, highlighting the effectiveness of our dual-field hemodynamic attention integration.

